# The causal role of transcranial alternating current stimulation at alpha frequency in boosting visual perceptual learning

**DOI:** 10.1101/2021.04.08.438912

**Authors:** Qing He, Xin-Yue Yang, Baoqi Gong, Keyan Bi, Fang Fang

## Abstract

Extensive training improves our ability to perceive visual contents around us, a phenomenon known as visual perceptual learning (VPL). Numerous studies have been conducted to understand the mechanisms of VPL, while the neural oscillatory mechanisms underpinning VPL has yet to be elucidated. To this end, we adopted transcranial alternating current stimulation (tACS), a neuromodulatory technique that can alter ongoing brain rhythms in a frequency-specific manner by applying external weak electric fields, to stimulate targeted cortical areas in human subjects while they performed an orientation discrimination learning task. Five groups of subjects undertook five daily training sessions to execute the task. Four groups received occipital tACS stimulation at 10 Hz (alpha band), 20 Hz (beta band), 40 Hz (gamma band), or sham 10 Hz (sham), and one group was stimulated at the sensorimotor regions by 10 Hz tACS. Compared with the sham stimulation, occipital tACS at 10 Hz, but not at 20 Hz or 40 Hz, increased both the learning rate and performance improvement. However, when 10 Hz tACS was delivered to the sensorimotor areas, the modulatory effects of tACS were absent, suggesting that tACS modulated the orientation discrimination learning in a frequency- and location-specific manner. Moreover, the tACS-induced enhancement lasted at least two months after the termination of training. Our findings provide strong evidence for the causal role of alpha oscillations in VPL and shed new light on the design of effective neuromodulation protocols that might facilitate rehabilitation for patients with neuro-ophthalmological disorders.

**Significance Statement:** Performance of visual tasks can be enhanced substantially by training, which is known as visual perceptual learning (VPL). However, little is known about the neural oscillatory mechanisms underlying VPL. To probe the causal link between a given oscillatory frequency band and VPL, transcranial alternating current stimulation (tACS) was applied while subjects performed an orientation discrimination learning task. Our results revealed that tACS modulates VPL in a frequency- and location-specific manner. Specifically, only training coupled with 10 Hz tACS over the occipital cortex speeded up the learning process and amplified the performance gain. Our findings demonstrate the causal role of alpha oscillations in VPL, and provide insight into developing more effective and efficient remediation protocols for clinical applications, e.g., amblyopia.

## Introduction

Repetitive visual experience or practice results in a dramatic and long-lasting improvement in perceiving visual contents, which is referred to as visual perceptual learning (VPL) (Maniglia & Seitz, 2018; Watanabe & Sasaki, 2015). VPL is usually viewed as a manifestation of experience-dependent neural plasticity in human brain (Lu *et al*., 2021). Numerous studies have been conducted to explore the neural substrates of VPL. It has been suggested that VPL occurs at multiple loci in the brain, from subcortical nuclei to high-level cortical areas involved in decision-making or attention (Bi *et al*., 2014; Gold *et al*., 2008; Mayhew *et al*., 2012; Mukai *et al*., 2007; Sanayei *et al*., 2018; Yu *et al*., 2016), and is manifested in various forms, such as enhanced neural response (Furmanski *et al*., 2004; Hua *et al*., 2010), refined neural representation (Chen *et al*., 2016; Jehee *et al*., 2012), sharpened tuning curve (Schoups *et al*., 2001; Yang & Maunsell, 2004), channel reweighting from sensory inputs to decision units (Chen *et al*., 2015; Dosher & Lu, 1998; Law & Gold, 2008; Yin Yan *et al*., 2014).

Neural oscillations, characterized by rhythmic changes in neural activity in a wide range of frequencies, play a critical role in cognitive functioning (Buzsaki & Draguhn, 2004; Thut *et al*., 2012). Aberrant neural oscillations might be associated with some neurological and psychiatric disorders (Schnitzler & Gross, 2005). However, the neural oscillatory mechanisms of VPL remain poorly understood and controversial. To date, only sparse studies have explored the neural oscillatory mechanisms of VPL, and results of these studies are not consistent. The focus of these studies is whether alpha- and/or gamma-band oscillations are associated with VPL. Human electroencephalogram (EEG) studies found that there was increased alpha power in parietal-occipital areas after training on visual tasks, such as visual search (Bays *et al*., 2015), perceptual grouping (Nikolaev *et al*., 2016), and orientation categorization (Muller-Gass *et al*., 2017). Pre-training resting-state alpha power could predict one’s learning performance (Muller-Gass *et al*., 2017). On the other hand, some studies showed that training induced an increase in gamma power (Gruber *et al*., 2001; Gruber *et al*., 2002; La Rocca *et al*., 2020), or changes in both alpha and gamma power (Hamame *et al*., 2011). In these studies, oscillatory neural activity was recorded before and after training, but was not manipulated during training to modulate VPL, which could not reveal the causal link between neural oscillations and VPL.

Transcranial alternating current stimulation (tACS) is a neuromodulation technique that can non-invasively alter brain excitability in a frequency-specific manner by delivering weak alternating currents to the scalp (Kanai *et al*., 2008; Kasten *et al*., 2019; Krause *et al*., 2019; Liu *et al*., 2018; Voroslakos *et al*., 2018). It is a wide-used tool for establishing a causal link between neural oscillations and cognition (Cabral-Calderin & Wilke, 2019; Johnson *et al*., 2020), across a broad range of frequencies and tasks (Antal *et al*., 2008; Helfrich *et al*., 2014b; Kar & Krekelberg, 2014; Pogosyan *et al*., 2009; Reinhart & Nguyen, 2019; Y. Zhang *et al*., 2019). However, tACS has not been applied to modulate VPL yet.

To this end, we stimulated human subjects’ visual cortex by using tACS at a given frequency (10 Hz, 20 Hz, or 40 Hz) while they were practicing an orientation discrimination task subserved by early visual cortex (Bang *et al*., 2018; Jia *et al*., 2020). We hypothesized that if neural oscillations of a given frequency band play an important role in the orientation discrimination learning, then applying tACS at that frequency band to the visual cortex will modulate the learning process and performance improvement.

## Materials and Methods

### Subjects

A total of 87 healthy subjects (58 females, 22.03 ± 3.96 years) participated in the present study. Subjects were assigned to one of five groups (Table 1): Group 1 received occipital tACS at 10 Hz (the 10 Hz occipital stimulation group), Group 2 received occipital tACS at 20 Hz (the 20 Hz occipital stimulation group), Group 3 received occipital tACS at 40 Hz (the 40 Hz occipital stimulation group), Group 4 received bilateral sensorimotor tACS at 10 Hz (the 10 Hz sensorimotor stimulation group), and Group 5 received occipital sham stimulation (the sham occipital stimulation group). The group sizes were determined by a power analysis (α = 0.05, two-tailed, power = 80%) based on our pilot study. A screening questionnaire was administrated before starting the study for each subject. Subjects were included if they met the following criteria: 1) were right-handed; 2) were aged between 18–40 years; 3) had normal or corrected-to-normal vision. Subjects were excluded if they met the following criteria: 1) had a history of neural surgery or epileptic seizures, any psychiatric or neurological disorders; 2) had sleep disorders or a total sleep time less than eight hours per night over the last two weeks; 3) were during the menstrual cycle or pregnancy (Fertonani *et al*., 2011; He *et al*., 2019). The present study was approved by the Ethics Committee of School of Psychological and Cognitive Sciences, Peking University. Written consent was obtained from each subject prior to the beginning of this study.

**Table1.** Demographic information of subjects

| Group | Sample size | Female/Male | Mean age (SD) | Age range |
| --- | --- | --- | --- | --- |
| Group 1 | 17 | 13/4 | 22.35 (5.24) | 18-37 |
| Group 2 | 17 | 11/6 | 21.47 (4.29) | 18-34 |
| Group 3 | 18 | 11/7 | 20.50 (2.07) | 18-24 |
| Group 4 | 18 | 12/6 | 23.83 (4.16) | 19-37 |
| Group 5 | 17 | 11/6 | 22.00 (2.98) | 18-28 |

### Apparatus and Stimulation protocol

Visual stimuli were generated and controlled using MATLAB (version 7.0) and Psychotoolbox-3 extensions (Brainard, 1997; Pelli, 1997) and were presented on a 19-inch Sony Trinitron color monitor (spatial resolution = 1600 × 1200 pixels, frame rate = 85 Hz) with a grey background (mean luminance = 47.59 cd/m^2^). The experiments were run in a dimly lit room. The monitor was the only source of light in the room. A chin-and-head rest was used to stabilize the head at a viewing distance of 65 cm. Subject’s eye movements were monitored by an Eyelink 1000 plus eye-tracking system (SR Research Ltd., Ontario, Canada) throughout the whole experiment.

Electrical stimulation was delivered using the DC-STIMULATOR MC (neuroConn GmbH, Ilmenau, Germany) through a pair of rubber electrodes of 5×5 cm^2^. The electrodes inserted in two soaked sponges (0.9% saline solution) were attached to the subjects’ scalp using elastic bandages. A sinusoidal current with a peak-to-peak intensity of 1.5 mA was administrated while subjects performed the training task. Both the DC offset and the phase difference between the two stimulation electrodes were set at zero. The impedance was constantly lower than 8 KΩ during all stimulation sessions.

### Visual stimulus and task

Oriented Gabor patches (diameter = 1.25°; spatial frequency = 3.0 cycle/°; Michelson contrast = 0.5; standard deviation of Gaussian envelope = 0.42°; random spatial phase) were located 5° from fixation in the lower left quadrant of the visual field. Gabor patches were spatially masked by noise using a pixel replacement method at a given signal-to-noise (S/N) ratio (Shibata *et al*., 2017). In our study, the S/N ratio was 75%, which meant that 25% of the pixels in the Gabor patch were replaced with random noise. In each trial, a small fixation point was displayed for 500 ms firstly, and then two Gabor patches with orientations of 26° and 26° + *θ* were presented successively for 100 ms each and were separated by a 500 ms blank interval (Figure 1(A)). These two Gabor patches were presented in random order. Subjects were instructed to make a two-alternative forced-choice (2AFC) judgment of the orientation of the second Gabor patch relative to the first one (clockwise or counter-clockwise) by pressing a key. The *θ* varied trial by trial and was adaptively controlled by a QUEST staircase to estimate subjects’ discrimination threshold at 75% correct (Watson & Pelli, 1983). Subjects were asked to have a rest with their eyes closed after each staircase. Feedback was not provided in all test and training sessions.

**Figure 1.**
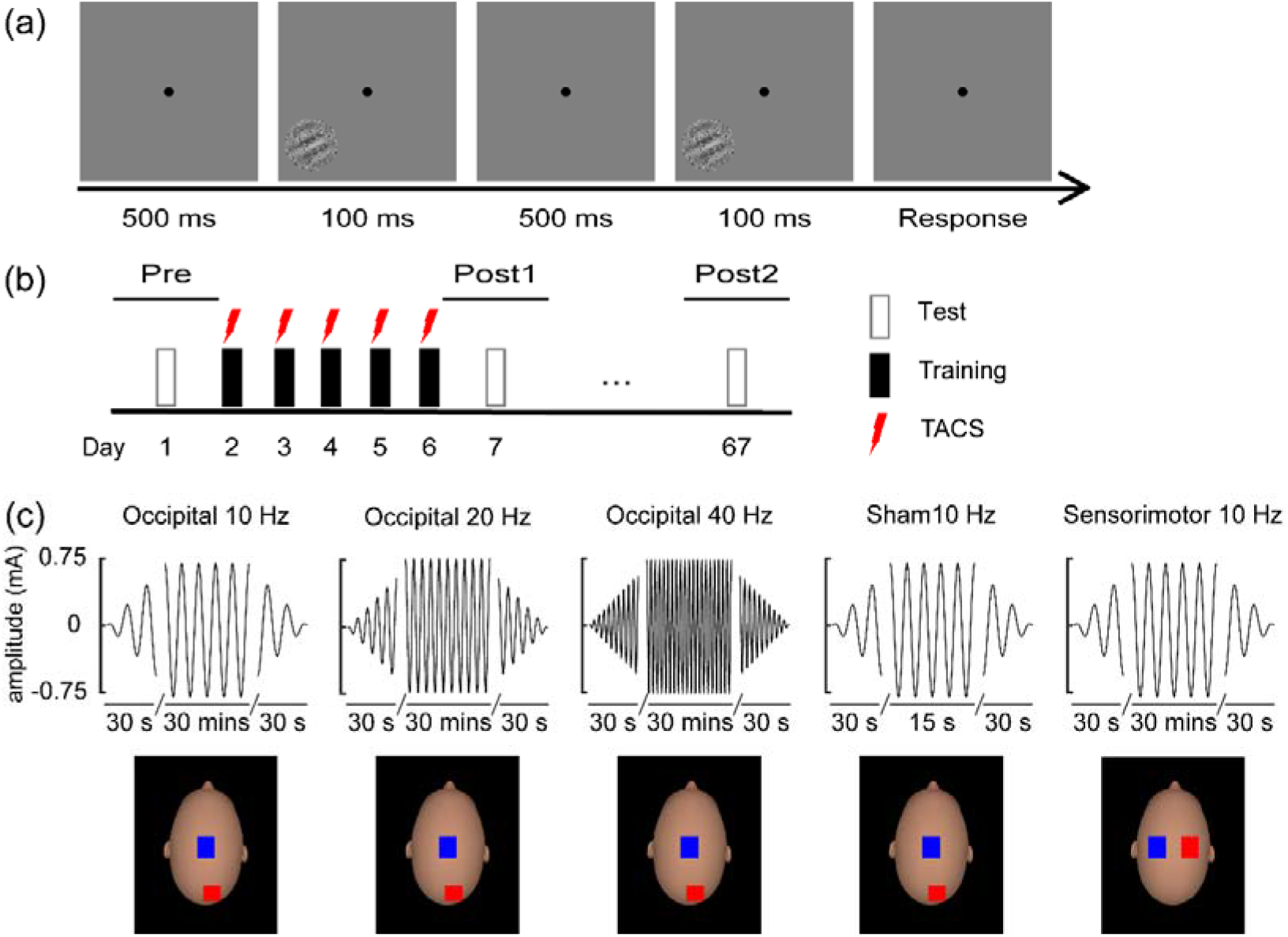
Stimuli, experimental protocol, and electrical stimulation protocol. (A) Schematic description of a 2AFC trial in a QUEST staircase for measuring orientation discrimination thresholds. Subjects were instructed to make a judgment of the orientation in the second interval relative to that in the first interval (clockwise or counter-clockwise), while gazing at the central fixation point. There was no feedback after each trial. (B) Experimental protocol. Subject underwent one pre-training test, five daily training sessions, and two post-training tests. The pre-training test (Pre) and post-training test 1 (Post1) and test 2 (Post2) took place on the days before, immediately after, and two months after training. The tACS was concurrently administrated during each training session. (C) Electrical stimulation protocol and montage. TACS at different frequencies (10 Hz, 20 Hz, and 40 Hz) were applied over different cortical regions. For occipital stimulation groups, stimulation electrodes were positioned over the occipital cortex (O2) and the vertex (Cz), while for the sensorimotor stimulation group, stimulation electrodes were positioned over the sensorimotor regions of both hemispheres. The electrode positions were identified by the international 10-20 EEG system. The red square and blue square denoted the anodal electrode and cathodal electrode, respectively. These heads were generated by FaceGen Modeller (version 3.4).

### Design

A single-blind, sham-controlled, between-subject design was adopted to explore the role of tACS in modulating the orientation discrimination learning. Subjects underwent five daily training sessions of the orientation discrimination task, which was preceded by a pre-training test (Pre) and was followed by two post-training tests . One of the post-training tests was completed immediately after training, i.e., Post1, and the other one (Post2) was conducted at least two months after Post1 (Figure 1(B)). Each participant completed six QUEST staircases of 50 trials at Pre, Post1, and Post2, and nine QUEST staircases during each training session.

First, to investigate which stimulation frequency is able to modulate the orientation discrimination learning, we recruited three groups of subjects to take part in this study, i.e., the 10 Hz, 20 Hz, and 40 Hz occipital stimulation groups. The choice of stimulation frequency of tACS was based on the controversy of underlying neural oscillatory mechanisms of VPL (Bays *et al*., 2015; Gruber *et al*., 2002; Hamame *et al*., 2011) and the study that found visual plasticity could be induced by 20 Hz tACS (Kanai *et al*., 2010). Also, 10 Hz, 20 Hz, and 40 Hz are representative frequencies of alpha band (8-12 Hz), beta band (13-30 Hz), and gamma band (above 30 Hz) (Helfrich *et al*., 2014a; Kanai *et al*., 2010; Nakazono *et al*., 2020; Struber *et al*., 2014; Wach *et al*., 2013; Zaehle *et al*., 2010). Two electrodes were placed over subjects’ visual cortex and the vertex (i.e., O2 and Cz in the international 10-20 EEG system), respectively (de Graaf *et al*., 2020; Fertonani *et al*., 2011; Hubel & Wiesel, 1968; Jehee *et al*., 2012; Jia *et al*., 2020) (Figure 1(C)). The stimulated hemisphere was contralateral to the visual field where the Gabor patches were presented. Subjects received electrical stimulation at a given frequency for about 27 minutes with a ramp-up of 30 s during each training session.

Second, we asked whether the observed modulatory effects by the 10 Hz tACS (see Results) were due to potential indirect effects, such as placebo effect. To this end, a fourth group of subjects were recruited to participate in our study, i.e., the sham occipital stimulation group. The electrical stimulation parameters were the same as those in the 10 Hz occipital stimulation group, except that subjects in the sham occipital stimulation group received the 10 Hz tACS for 15 seconds with a ramp up/down of 30 s at the beginning of each training session.

Finally, to exam whether the 10 Hz tACS modulated the orientation discrimination learning in a stimulation location-specific manner, a fifth group of subjects were recruited, i.e., the 10 Hz sensorimotor stimulation group. The stimulation setting was similar to that in the 10 Hz occipital stimulation group, except that two electrodes were positioned over the bilateral sensorimotor cortical regions (Cappelletti *et al*., 2013; Cappelletti *et al*., 2015), i.e., C1 and C2.

### Statistical analysis methods

For each test or training session, the estimated threshold was defined as the geometric mean of thresholds from all QUEST staircases. Percent improvement, which describes changes in performance after training, was calculated as (*pre-training threshold* − *post-training threshold*)/*pre-training threshold* × *100%*. To illustrate the threshold dynamics during the training course, a power function was used to fit the learning curves of orientation discrimination across all test and training sessions (Jeter *et al*., 2009; P. Zhang *et al*., 2018):

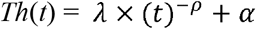

Where *Th* is the predicted orientation discrimination threshold, *t* is the number of training sessions, λ is the initial threshold, ρ is the learning rate, and *α* denotes the minimum threshold achieved after training. A nonlinear least-square method, implemented in MATLAB (MathWorks, Natick, MA), was used to minimize the sum of squared differences between model predictions and observed values.

Orientation discrimination thresholds were analyzed using a mixed-design analysis of variance (ANOVA) with a between-subjects factor *Group* (occipital 10 Hz, 20 Hz, 40 Hz, sham, and sensorimotor 10 Hz) and a within-subjects factor *Test* (Pre, Post1 and Post2). Learning rates and percent improvements were analyzed using ANOVA with a between-subjects factor of *Group*. For multiple comparisons, Benjamini-Hochberg method (BH) was used to control false discovery rate (FDR)(Benjamini & Hochberg, 1995; Benjamini & Yekutieli, 2001). For ANOVAs, 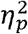 was computed as a measure of effect size. For t-tests, Cohen’s d was computed as a measure of effect size. Statistical analyses were conducted using R (R Core Team, 2020).

## Results

### Occipital tACS modulated orientation discrimination learning in a frequency-specific manner

First, subjects stimulated by occipital tACS at 10 Hz, 20 Hz, and 40 Hz started with comparable performance at Pre (*F*(2, 49) = 0.33, *p* = 0.72, 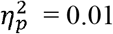). As shown in Figure 2 (A), subjects’ orientation discrimination thresholds declined with training for all the training groups. For each subject, the orientation discrimination thresholds across all test and training sessions were fitted with a power function. A one-way ANOVA with *Group* (10 Hz, 20 Hz, and 40 Hz) as a between-subjects factor revealed that there were significant group differences in learning rate, i.e., the estimated ρ-value of the power function (*F*(2, 49) = 7.38, *p* = 0.002, 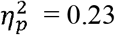).

**Figure 2.**
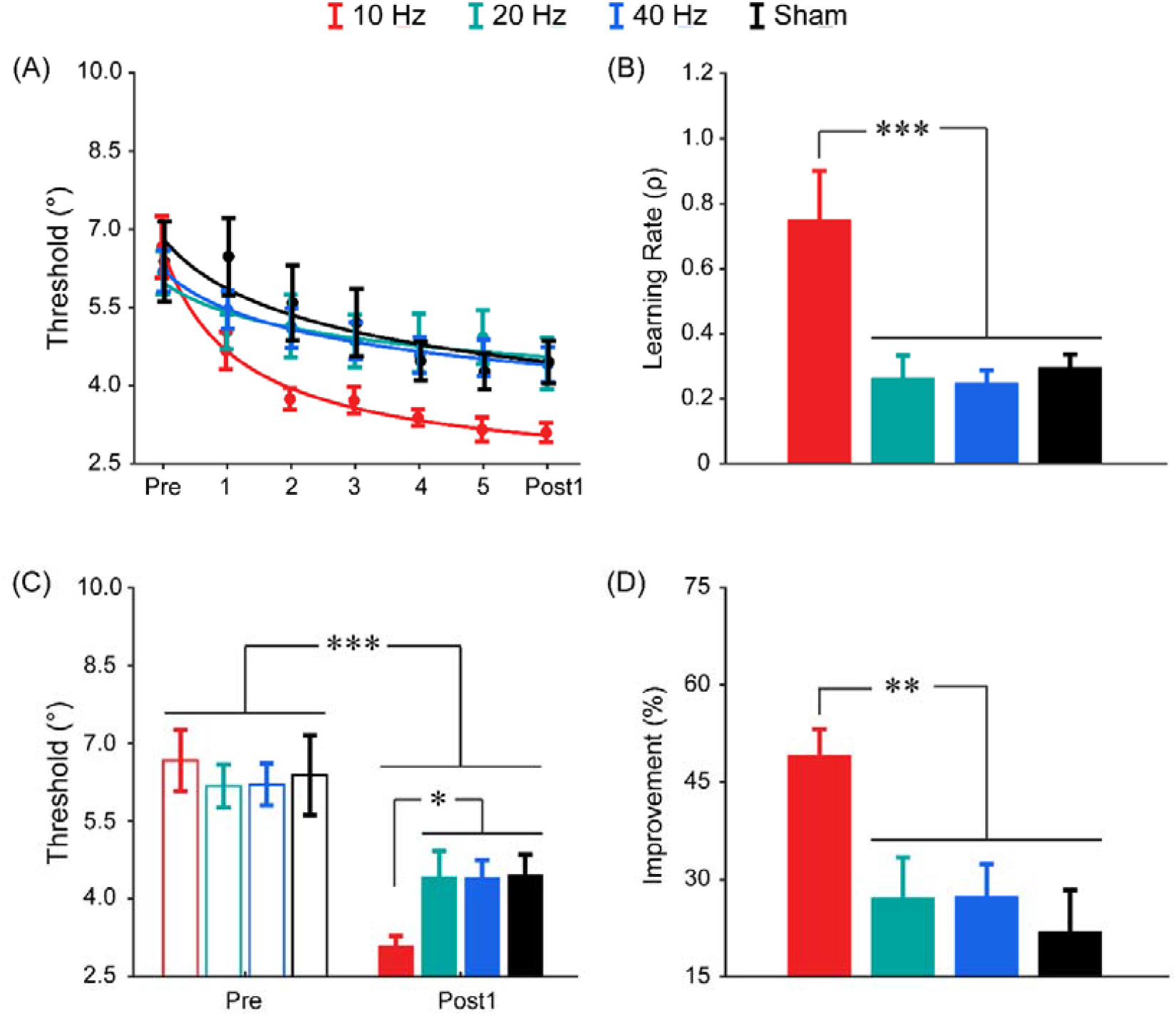
Results of the main experiment in which visual cortex was stimulated using tACS at different frequencies. (A) Learning curves. Dots represent averaged thresholds across subjects at different test and training days, and lines represent fitted learning curves using a power function. (B) Learning rates. (C) Orientation discrimination thresholds at Pre and Post1. (D) Percent improvements in orientation discrimination performance at Post1, relative to Pre. \**p* <.05; \*\**p* < .01; \*\*\**p* < .001. Error bars denote the standard error of the mean (SEM) across subjects.

*Post-hoc* analysis showed that the learning rate of the 10 Hz occipital stimulation group was significantly greater than those of the other two groups (10 Hz vs. 20 Hz: *t*(32) = 3.37, *p*_adj_ = 0.004, Cohen’s d =0.98; 10 Hz vs. 40 Hz: *t*(33) = 3.30, *p*_adj_ = 0.004, Cohen’s d = 1.09), and there was no significant difference in learning rate between the 20 Hz occipital stimulation and 40 Hz occipital stimulation groups (*t*(33) = -0.124, *p*_adj_ = 0.90) (Figure 2(B)). These results demonstrated that, relative to beta and gamma frequencies, tACS at alpha frequency accelerated the learning process.

Then subjects’ orientation discrimination thresholds were submitted to a mixed-design ANOVA with *Test* (Pre and Post1) as a within-subjects factor and *Group* (10 Hz, 20Hz, and 40 Hz) as a between-subjects factor to examine differences in performance. We found a significant main effect of *Test* (*F*(1, 49) = 83.48, *p* = 3.69 × 10^−12^, 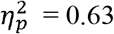), and a significant interaction between *Test* and *Group* (*F*(2, 49) = 5.32, *p* = 0.008, 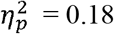), but no significant main effect of *Group* (*F*(2, 49) = 0.46, *p* = 0.63, 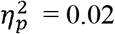). Further analyses showed that the thresholds at Post1 were lower than that those at Pre for all groups (all *t*s < -3.78, all *p*_adj_s < 0.002, and all Cohen’s ds > 0.92, by paired *t*-test), demonstrating significant learning effects occurred in an unsupervised manner here because no feedback was provided during learning (Loewenstein *et al*., 2021; Tsodyks & Gilbert, 2004; Weiss *et al*., 1993). At Post1, the threshold in the 10 Hz occipital stimulation group was significantly lower than those in the 20 Hz occipital stimulation group (*t*(32) = -2.58, *p*_adj_ = 0.039, Cohen’s d = 0.87) and the 40 Hz occipital stimulation group (*t*(33) = -2.57, *p*_adj_ = 0.039, Cohen’s d = 1.13). The threshold difference between the 20 Hz and 40 Hz occipital stimulation groups was not significant (*t*(33) = 0.05, *p* = 0.96) (Figure 2 (C)).

The above results suggested that there was greater performance improvement in the 10 Hz occipital stimulation group. To confirm this, we performed a one-way ANOVA on percent improvement. The statistical results showed that the differences in percent improvement among the three stimulation groups s were significant (*F*(2, 49) = 5.82, *p* = 0.005, 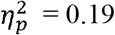). *Post-hoc* analysis revealed that the percent improvement in the 10 Hz occipital stimulation group was higher than those in the 20 Hz occipital stimulation group (*t*(32) = 2.96, *p*_adj_ = 0.014, Cohen’s d = 1.00) and the 40 Hz occipital stimulation group ((*t*(33) = 2.97, *p*_adj_ = 0.014, Cohen’s d = 1.13), and the difference between the 20 Hz and the 40 Hz occipital stimulation groups was not significant (*t*(33) = 0.03, *p* = 0.98) (Figure 2 (D)). These results demonstrated that, relative to beta and gamma frequencies, tACS at alpha frequency enabled subjects to acquire a greater improvement in the learning task.

To examine whether the observed modulatory effects from the 10 Hz tACS were due to possible indirect effects (e.g., placebo effect), one more group of subjects received the sham occipital stimulation. Note that the learning effect in the sham occipital stimulation group is presumably equivalent to the learning effect without electrical stimulation. Compared with the sham occipital stimulation group, only the 10 Hz occipital stimulation group showed a faster learning rate (*t*(32) = 3.79, *p*_adj_ = 0.002, Cohen’s d = 1.13), a lower threshold at Post1 (*t*(32) = -2.58, *p*_adj_ = 0.02, Cohen’s d = 1.06), and a greater improvement (*t*(32) = 3.48, *p*_adj_ = 0.005, Cohen’s d = 1.24), and these beneficial effects were absent in the 20 Hz and 40 Hz occipital stimulation groups (all *p*s > 0.90).

Previous studies have shown that the after-effects of tACS on behavioral performance and neural activity are temporally short (Clancy *et al*., 2018; Kasten *et al*., 2016; Neuling *et al*., 2013; Struber *et al*., 2015), while one remarkable characteristics of VPL is the long-term persistence of learning effects after the termination of training (Bi *et al*., 2014; Frank *et al*., 2020; P. Zhang *et al*., 2018). To examine whether the facilitatory effects on the orientation discrimination learning by the 10 Hz tACS is long-lasting, subjects in the 10 Hz occipital stimulation group and the sham occipital stimulation group were retested at least two months after Post1 (i.e., Post2). For subjects in the 10 Hz occipital stimulation group, their thresholds at Post2 were not significantly different from those at Post1 (*t*(16) = -0.37, *p* = 0.72). Similarly, there was no significant difference in threshold between Post1 and Post2 in the sham occipital stimulation group (*t*(16) = -5.2 × 10^−3^, *p* = 0.99) (Figure 3 (A)). Notably, some subjects (N = 9 and N = 12 in the 10 Hz occipital stimulation group and the sham occipital stimulation group, respectively) were retested at least 14 months after training, i.e., Post3. There was no significant difference in threshold between Post1 and Post3 (10 Hz: *t*(8) = 1.51, *p* = 0.17; sham: *t*(11) = 0.76, *p* = 0.47) (Figure 3 (B)). Therefore, the modulatory effect of 10 Hz tACS on the orientation discrimination learning was remarkably long-lasting.

**Figure 3.**
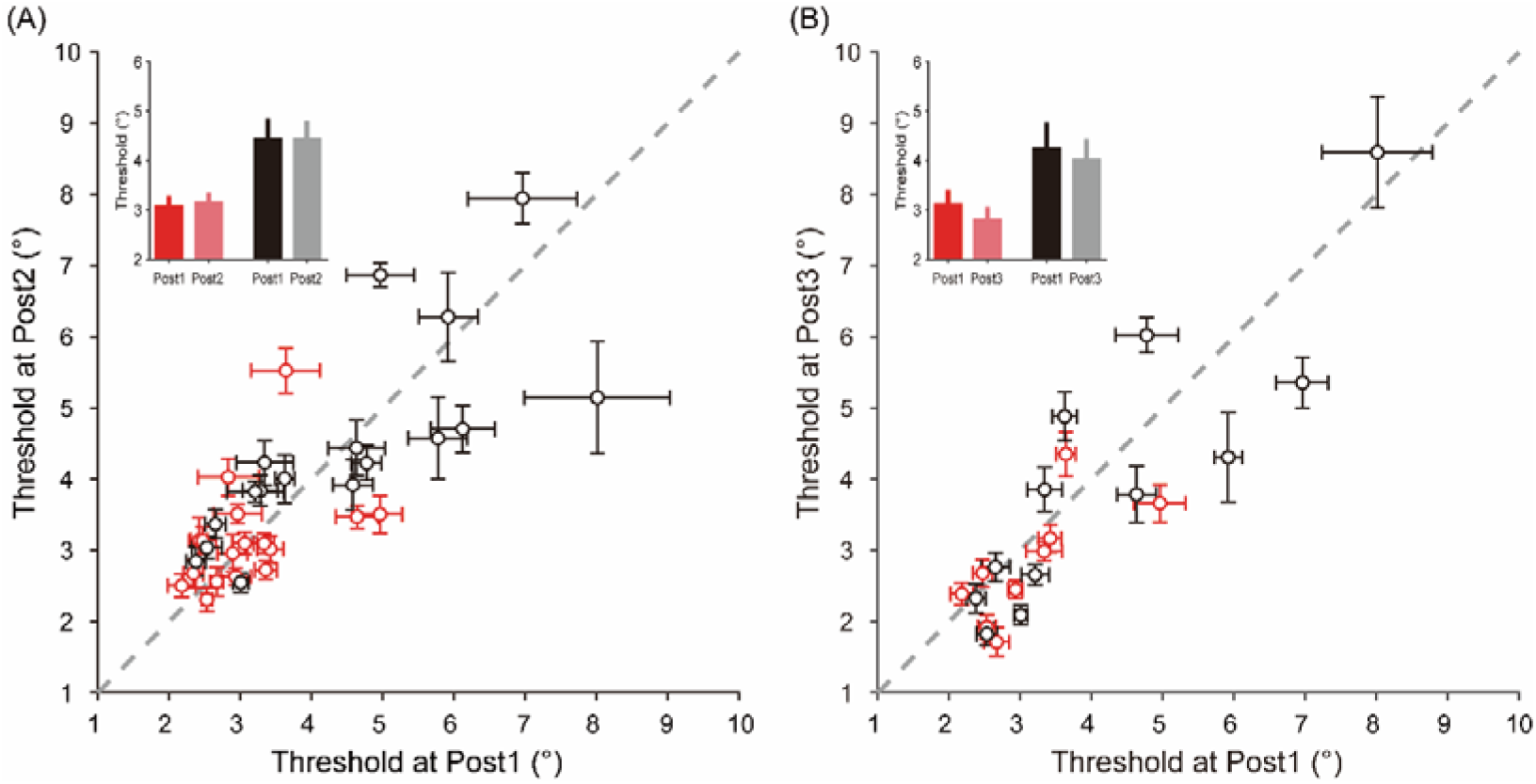
Retention of the learning effect. (A) Retest results two months after training for subjects in the 10 Hz occipital stimulation group and the sham occipital stimulation group. (B) Retest results 14 months after training for part of the subject cohort. Each red dot and black dot represent a single subject in the 10 Hz occipital stimulation group and the sham occipital stimulation group, respectively. The insets depict the mean thresholds for the 10 Hz occipital stimulation group (red and pink bars) and the sham occipital stimulation group (black and grey bars). Error bars denote the standard error of the mean (SEM) across QUEST staircases or subjects.

### 10 Hz tACS modulated orientation discrimination learning in a location-specific manner

To explore whether the 10 Hz tACS modulated the orientation discrimination learning in a location-specific manner, one more group of subjects (i.e., the 10 Hz sensorimotor stimulation group) were trained on the orientation discrimination task while their bilateral sensorimotor regions were stimulated by the 10 Hz tACS. For the three groups of subjects (the 10 Hz occipital stimulation group, the 10 Hz sensorimotor stimulation group, and the sham occipital stimulation group), they had about equal thresholds at Pre (*F*(2, 49) = 0.12, *p* = 0.88, 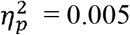). Extensive training reduced subjects’ orientation discrimination thresholds gradually for all groups (Figure 4(A)). Again, there were significant differences in learning rate among the three groups (*F*(2, 49) = 10.60, *p* = 1.49 × 10^−4^, 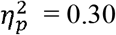). The learning rate of the 10 Hz occipital stimulation group was greater than that of the 10 Hz sensorimotor stimulation group (*t*(32) = 4.21, *p*_adj_ = 3.26 × 10^−4^, Cohen’s d = 1.27), but the difference in learning rate between the 10 Hz sensorimotor stimulation group and the sham occipital stimulation group was not significant (*t*(33) = -0.41, *p* = 0.68) (Figure 4(B)). These results on learning rate demonstrated that the orientation discrimination learning efficiency was not modulated by the 10 Hz tACS applied at sensorimotor areas.

**Figure 4.**
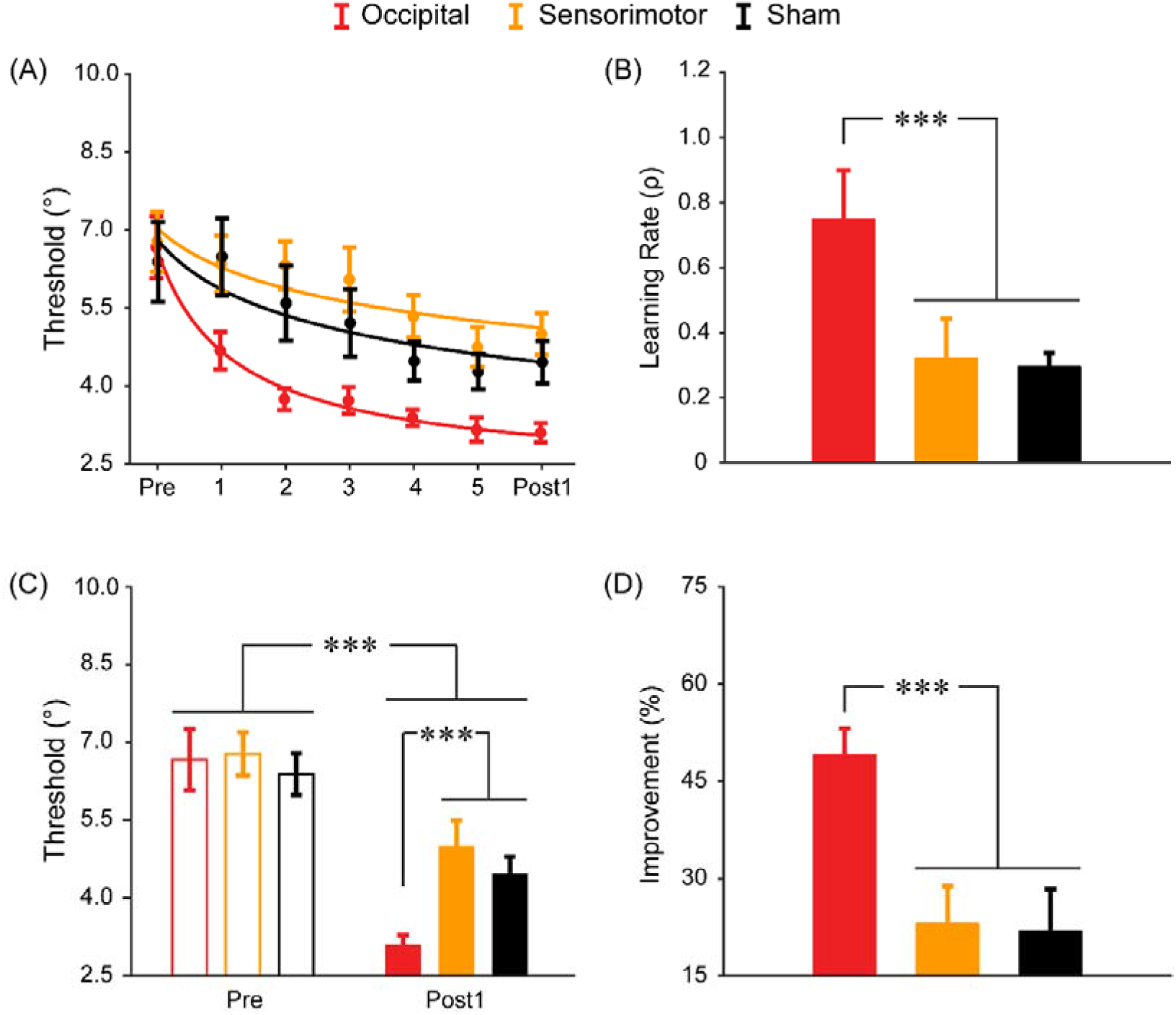
Results of the control experiment in which the 10 Hz tACS was applied to bilateral sensorimotor areas. (A) Learning curves. Dots represent averaged thresholds across subjects at different test and training days, and lines represent fitted learning curves using a power function. (B) Learning rates. (C) Orientation discrimination thresholds at Pre and Post1. (D) Percent improvements in orientation discrimination performance at Post1, relative to Pre . \*\*\**p* < .001. Error bars denote the standard error of the mean (SEM) across subjects.

Regarding orientation discrimination thresholds, a 2 × 3 mixed-design ANOVA with a within-subjects factor of *Test* (Pre and Post1) and a between-subjects factor of *Group* (occipital, sensorimotor, and sham) showed that the main effect of *Test* (*F*(1, 49) = 63.04, *p* = 2.36 × 10^−10^, 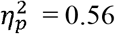) and the interaction between *Test* and *Group* (*F*(2, 49) = 4.33, *p* = 0.019, 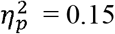) were significant, but the main effect of *Group* (*F*(2, 48) = 1.77, *p* = 0.18) was not. Paired *t*-test showed that the thresholds at Post1 were lower than those at Pre for all the three groups (all *t*s ≤-3.44, all *p*_adj_s ≤ 0.003, and all Cohen’s ds ≥ 0.83). A simple main effect analysis revealed that, at Post1, the threshold in the 10 Hz occipital stimulation group was lower than that in the 10 Hz sensorimotor stimulation group (*t*(33) = -4.94, *p*_adj_ = 1.50 × 10^−4^, Cohen’s d = 1.64), but the threshold difference between the 10 Hz sensorimotor stimulation group and the sham occipital stimulation group was not significant (*t*(33) = 1.43, *p* = 0.16) (Figure 4 (C)). Accordingly, there were significant differences in percent improvement between the three groups (*F*(2, 49) = 8.69, *p* = 5.88 × 10^−4^, 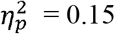). *Post-hoc* analysis showed that the percent improvement in the 10 Hz occipital stimulation group was higher than that in the 10 Hz sensorimotor stimulation group (*t*(33) = 4.13, *p*_adj_ = 0.0013, Cohen’s d = 1.38), while there was no significant difference between the 10 Hz sensorimotor stimulation group and the sham occipital stimulation group (*t*(33) = -0.25, *p* = 0.81) (Figure 4(D)). These results demonstrated that the 10 Hz tACS delivered over sensorimotor areas did not result in more performance improvements in the orientation discrimination learning.

## Discussion

To better understand the neural oscillatory mechanisms underlying VPL, tACS was administered while subjects were undertaking an orientation discrimination learning task. Our study showed that, relative to sham stimulation, 10 Hz tACS applied to visual cortex accelerated learning and led to greater improvement in task performance. The facilitatory effects were absent when visual cortex was stimulated by tACS at other frequencies (20 Hz and 40 Hz) or when other cortical areas (sensorimotor cortical areas) were stimulated by 10 Hz tACS, indicating that tACS modulated VPL in a frequency- and location-specific manner. To the best of our knowledge, this is the first study demonstrating the effects of tACS on VPL. Our results provide strong evidence of the causal role of occipital alpha band oscillations in VPL, updating our knowledge on the role of alpha oscillations in neural plasticity. Moreover, it is also the first study showing that the enhanced performance by combining perceptual training and non-invasive brain stimulation could last at least one year after training, which provides a promising and feasible way for clinical applications.

Our finding of the causal role of occipital alpha oscillations in VPL is consistent with previous human EEG studies across different visual learning tasks (Bays *et al*., 2015; Muller-Gass *et al*., 2017; Nikolaev *et al*., 2016; Toosi *et al*., 2017), and is also consistent with tactile perceptual learning studies (Brickwedde *et al*., 2019; Freyer *et al*., 2013; Freyer *et al*., 2012). These EEG learning studies suggested that the alpha power enhancement in parietal-occipital regions is a manifestation of the neural mechanisms of perceptual learning. Furthermore, both the resting-state alpha power before learning (Freyer *et al*., 2013; Muller-Gass *et al*., 2017) and the learning-induced alpha power changes during task execution (Brickwedde *et al*., 2019; Freyer *et al*., 2013) correlated with individual’s learning outcomes, demonstrating a close association between alpha oscillations and perceptual learning. Taken together, all these studies point to a pivotal role of alpha oscillations in perceptual learning.

We note that our study is different from some previous studies in scientific finding and method. First, several human EEG studies found that training induced increases in gamma or beta band oscillations (Gruber *et al*., 2001; Gruber *et al*., 2002; Hamame *et al*., 2011; La Rocca *et al*., 2020; Theves *et al*., 2020). Several factors could explain this discrepancy. For instance, different stimuli and tasks, which engaged distinctive processing mechanisms, were used in these learning studies (Brunet & Fries, 2019; Gruber *et al*., 2002). Also, in these studies, EEG was recorded before and after training, while tACS was administrated during training in our study. Second, previous studies found that transcranial direct current stimulation (tDCS) and transcranial random noise stimulation (tRNS) were able to modulate VPL (Cappelletti *et al*., 2013; Contemori *et al*., 2019; Fertonani *et al*., 2011; Frangou *et al*., 2018; Herpich *et al*., 2019; Van Meel *et al*., 2016). However, both tDCS and tRNS, with less understood working mechanisms, cannot alter brain oscillations in a frequency-specific manner, which limits our understanding of the neural oscillatory mechanisms of VPL.

Previous studies on the biophysical mechanisms of tACS have already provided strong support for the efficacy of tACS on altering neural oscillations. TACS applied over the scalp can generate changes in intracerebral electric fields in non-human primates (Johnson *et al*., 2020; Kar *et al*., 2017; Krause *et al*., 2019; Vieira *et al*., 2020) and humans (Opitz *et al*., 2016; Voroslakos *et al*., 2018), thereby modifying ongoing neural oscillations in the targeted brain region (Helfrich *et al*., 2014b; Kar *et al*., 2017; Kasten *et al*., 2018; Vossen *et al*., 2015; Zaehle *et al*., 2010). The tACS effects are usually manifested as increased neural spectral power and/or phase synchronization (Helfrich *et al*., 2014b; Huang *et al*., 2021), which is confirmed by our EEG experiment. In this experiment, we recorded subjects’ resting-state EEG signals for 10 minutes both before and after 25-min 10Hz tACS over O2, which is roughly equivalent to the stimulation time in the main experiment. We analyzed 1-min EEG signals before and after the stimulation from Oz, O2, POz, and PO4 (Kasten *et al*., 2016). We found that subjects’ resting-state alpha power after the tACS stimulation was significantly higher than that before the stimulation. It is noteworthy that, in our study, tACS at alpha frequency modulated VPL in a location-specific manner. This ruled out possibly indirect effects, such as retinal stimulation, transcutaneous stimulation of peripheral nerves, and placebo effects (Asamoah *et al*., 2019; Liu *et al*., 2018; Turi *et al*., 2013). Taken together, these results suggest that occipital tACS at 10 Hz entrained endogenous alpha oscillations in visual cortex and thereby facilitated VPL causally.

The functional role of alpha band neural oscillations in visual perception, attention, and other cognitive functions has been recognized for a long time. Rather than only reflecting an idling state of the brain (Clayton *et al*., 2018; Sigala *et al*., 2014), it functions to gate ongoing sensory information processing (Jensen & Mazaheri, 2010; VanRullen, 2016). According to a dominant view about the function of alpha oscillations - the inhibition hypothesis, alpha oscillations are assumed to actively suppress processing of irrelevant sensory information and therefore direct computational neural resources to task-relevant events of higher priority (Klimesch *et al*., 2007). Specifically, alpha power is correlated with neural excitability negatively (Lange *et al*., 2013; Romei *et al*., 2008), and higher alpha power is predictive of decreased visual detection and discrimination performance (Ergenoglu *et al*., 2004; Mathewson *et al*., 2014; Roberts *et al*., 2014; van Dijk *et al*., 2008; Zazio *et al*., 2020). Since many studies have demonstrated that alpha tACS is able to increase ongoing alpha power (Clayton *et al*., 2019; Helfrich *et al*., 2014b; Kasten *et al*., 2019; Nakazono *et al*., 2020; Zaehle *et al*., 2010; Y. Zhang *et al*., 2019), the inhibition hypothesis cannot explain our finding here straightforwardly. Notably, it has been argued that alpha oscillations also reflect cortical feedback mechanisms in visual cortex (Keller *et al*., 2020; Mejias *et al*., 2016; Michalareas *et al*., 2016; van Kerkoerle *et al*., 2014). For example, using intracranial multi-array recording in monkeys, researchers found that neural activity in primary visual area V1 was driven by feedback signals in the alpha band from visual area V4 (Bastos *et al*., 2015; van Kerkoerle *et al*., 2014). This interregional feedback projections from V4 to V1 was also confirmed in human magnetoencephalography (MEG) study (Michalareas *et al*., 2016). Recently, studies from several research groups proposed that VPL could be implemented, at least partially, by cortical feedback connections that propagated neural signals from higher to lower areas (Crist *et al*., 2001; Jia *et al*., 2020; Moldakarimov *et al*., 2014; Mukai *et al*., 2007; Schafer *et al*., 2007; Y. Yan *et al*., 2018). Under this framework, entrained alpha oscillations by 10 Hz tACS might boost VPL by strengthening cortico-cortical feedback connections, which can be investigated in the future

Our findings not only deepen our understanding of the neural mechanisms underlying VPL, but also offer a promising guide for future clinical intervention to impaired visual functions. In clinical practice, efficacy/effectiveness, efficiency, and persistence are three of the most remarkable factors that constrain the application of perceptual enhancement methods or techniques (Herpich *et al*., 2019; Rodan *et al*., 2020). In our study, we found that applying tACS at 10 Hz over the visual cortex is able to help observers obtain more benefits within a shorter time. What is more, the modulatory effect of occipital 10 Hz tACS on visual performance was able to last for a very long period. Therefore, our training and stimulation protocol can be used as a potential treatment to help rehabilitate impaired visual functions in clinical populations, such as, patients with neuro-ophthalmological disorders (Levi & Li, 2009).

## Acknowledgements

We thank Rui-Ling Cai for her help in data collection, and Dr. Qian Wang for her helpful comments on the draft. This work was supported by the National Natural Science Foundation of China (31930053), Beijing Municipal Science and Technology Commission (Z181100001518002) and Beijing Academy of Artificial Intelligence (BAAI).

## Reference

Antal, A., Boros, K., Poreisz, C., Chaieb, L., Terney, D., & Paulus, W. (2008). Comparatively weak after-effects of transcranial alternating current stimulation (tACS) on cortical excitability in humans. Brain Stimul, 1(2), 97–105.

Asamoah, B., Khatoun, A., & Mc Laughlin, M. (2019). tACS motor system effects can be caused by transcutaneous stimulation of peripheral nerves. Nat Commun, 10(1), 266.

Bang, J. W., Sasaki, Y., Watanabe, T., & Rahnev, D. (2018). Feature-Specific Awake Reactivation in Human V1 after Visual Training. J Neurosci, 38(45), 9648–9657.

Bastos, A. M., Litvak, V., Moran, R., Bosman, C. A., Fries, P., & Friston, K. J. (2015). A DCM study of spectral asymmetries in feedforward and feedback connections between visual areas V1 and V4 in the monkey. Neuroimage, 108, 460–475.

Bays, B. C., Visscher, K. M., Le Dantec, C. C., & Seitz, A. R. (2015). Alpha-band EEG activity in perceptual learning. J Vis, 15(10), 1–12.

Benjamini, Y., & Hochberg, Y. (1995). Controlling the False Discovery Rate: A Practical and Powerful Approach to Multiple Testing. J. R. Stat. Soc. B, 57(1), 289–300.

Benjamini, Y., & Yekutieli, D. (2001). The control of the false discovery rate in multiple testing under dependency. Ann. Statist., 29(4), 1165–1188.

Bi, T., Chen, J., Zhou, T., He, Y., & Fang, F. (2014). Function and structure of human left fusiform cortex are closely associated with perceptual learning of faces. Curr Biol, 24(2), 222–227.

Brainard, D. H. (1997). The Psychophysics Toolbox. Spat Vis, 10(4), 433–436.

Brickwedde, M., Kruger, M. C., & Dinse, H. R. (2019). Somatosensory alpha oscillations gate perceptual learning efficiency. Nat Commun, 10(1), 263.

Brunet, N. M., & Fries, P. (2019). Human visual cortical gamma reflects natural image structure. Neuroimage, 200, 635–643.

Buzsaki, G., & Draguhn, A. (2004). Neuronal oscillations in cortical networks. Science, 304(5679), 1926–1929.

Cabral-Calderin, Y., & Wilke, M. (2019). Probing the Link Between Perception and Oscillations: Lessons from Transcranial Alternating Current Stimulation. Neuroscientist, 26(1), 57–73.

Cappelletti, M., Gessaroli, E., Hithersay, R., Mitolo, M., Didino, D., Kanai, R., … Walsh, V. (2013). Transfer of cognitive training across magnitude dimensions achieved with concurrent brain stimulation of the parietal lobe. J Neurosci, 33(37), 14899–14907.

Cappelletti, M., Pikkat, H., Upstill, E., Speekenbrink, M., & Walsh, V. (2015). Learning to integrate versus inhibiting information is modulated by age. J Neurosci, 35(5), 2213–2225.

Chen, N., Bi, T., Zhou, T., Li, S., Liu, Z., & Fang, F. (2015). Sharpened cortical tuning and enhanced cortico-cortical communication contribute to the long-term neural mechanisms of visual motion perceptual learning. Neuroimage, 115, 17–29.

Chen, N., Cai, P., Zhou, T., Thompson, B., & Fang, F. (2016). Perceptual learning modifies the functional specializations of visual cortical areas. Proc Natl Acad Sci U S A, 113(20), 5724–5729.

Clancy, K. J., Baisley, S. K., Albizu, A., Kartvelishvili, N., Ding, M., & Li, W. (2018). Lasting connectivity increase and anxiety reduction via transcranial alternating current stimulation. Soc Cogn Affect Neurosci, 13(12), 1305–1316.

Clayton, M. S., Yeung, N., & Cohen Kadosh, R. (2018). The many characters of visual alpha oscillations. Eur J Neurosci, 48(7), 2498–2508.

Clayton, M. S., Yeung, N., & Cohen Kadosh, R. (2019). Electrical stimulation of alpha oscillations stabilizes performance on visual attention tasks. J Exp Psychol Gen, 148(2), 203–220.

Contemori, G., Trotter, Y., Cottereau, B. R., & Maniglia, M. (2019). tRNS boosts perceptual learning in peripheral vision. Neuropsychologia, 125, 129–136.

Crist, R. E., Li, W., & Gilbert, C. D. (2001). Learning to see: experience and attention in primary visual cortex. Nat Neurosci, 4(5), 519–525.

de Graaf, T. A., Thomson, A., Janssens, S. E. W., van Bree, S., Ten Oever, S., & Sack, A. T. (2020). Does alpha phase modulate visual target detection? Three experiments with tACS-phase-based stimulus presentation. Eur J Neurosci, 51(11), 2299–2313.

Dosher, B. A., & Lu, Z. L. (1998). Perceptual learning reflects external noise filtering and internal noise reduction through channel reweighting. Proc Natl Acad Sci U S A, 95(23), 13988–13993.

Ergenoglu, T., Demiralp, T., Bayraktaroglu, Z., Ergen, M., Beydagi, H., & Uresin, Y. (2004). Alpha rhythm of the EEG modulates visual detection performance in humans. Brain Res Cogn Brain Res, 20(3), 376–383.

Fertonani, A., Pirulli, C., & Miniussi, C. (2011). Random noise stimulation improves neuroplasticity in perceptual learning. J Neurosci, 31(43), 15416–15423.

Frangou, P., Correia, M., & Kourtzi, Z. (2018). GABA, not BOLD, reveals dissociable learning-dependent plasticity mechanisms in the human brain. eLife, 7, e35854.

Frank, S. M., Qi, A., Ravasio, D., Sasaki, Y., Rosen, E. L., & Watanabe, T. (2020). Supervised Learning Occurs in Visual Perceptual Learning of Complex Natural Images. Curr Biol, 30(15), 2995–3000 e2993.

Freyer, F., Becker, R., Dinse, H. R., & Ritter, P. (2013). State-dependent perceptual learning. J Neurosci, 33(7), 2900–2907.

Freyer, F., Reinacher, M., Nolte, G., Dinse, H. R., & Ritter, P. (2012). Repetitive tactile stimulation changes resting-state functional connectivity-implications for treatment of sensorimotor decline. Front Hum Neurosci, 6, 144.

Furmanski, C. S., Schluppeck, D., & Engel, S. A. (2004). Learning strengthens the response of primary visual cortex to simple patterns. Curr Biol, 14(7), 573–578.

Gold, J. I., Law, C. T., Connolly, P., & Bennur, S. (2008). The relative influences of priors and sensory evidence on an oculomotor decision variable during perceptual learning. J Neurophysiol, 100(5), 2653–2668.

Gruber, T., Keil, A., & Muller, M. M. (2001). Modulation of induced gamma band responses and phase synchrony in a paired associate learning task in the human EEG. Neurosci Lett, 316(1), 29–32.

Gruber, T., Muller, M. M., & Keil, A. (2002). Modulation of induced gamma band responses in a perceptual learning task in the human EEG. J Cogn Neurosci, 14(5), 732–744.

Hamame, C. M., Cosmelli, D., Henriquez, R., & Aboitiz, F. (2011). Neural mechanisms of human perceptual learning: electrophysiological evidence for a two-stage process. PLoS One, 6(4), e19221.

He, Q., Lin, B. R., Zhao, J., Shi, Y. Z., Yan, F. F., & Huang, C. B. (2019). No effects of anodal transcranial direct current stimulation on contrast sensitivity function. Restor Neurol Neurosci, 37(2), 109–118.

Helfrich, R. F., Knepper, H., Nolte, G., Struber, D., Rach, S., Herrmann, C. S., … Engel, A. K. (2014a). Selective modulation of interhemispheric functional connectivity by HD-tACS shapes perception. PLoS Biol, 12(12), e1002031.

Helfrich, R. F., Schneider, T. R., Rach, S., Trautmann-Lengsfeld, S. A., Engel, A. K., & Herrmann, C. S. (2014b). Entrainment of brain oscillations by transcranial alternating current stimulation. Curr Biol, 24(3), 333–339.

Herpich, F., Melnick, M. D., Agosta, S., Huxlin, K. R., Tadin, D., & Battelli, L. (2019). Boosting Learning Efficacy with Noninvasive Brain Stimulation in Intact and Brain-Damaged Humans. J Neurosci, 39(28), 5551–5561.

Hua, T., Bao, P., Huang, C. B., Wang, Z., Xu, J., Zhou, Y., & Lu, Z. L. (2010). Perceptual learning improves contrast sensitivity of V1 neurons in cats. Curr Biol, 20(10), 887–894.

Huang, W. A., Stitt, I. M., Negahbani, E., Passey, D. J., Ahn, S., Davey, M., … Frohlich, F. (2021). Transcranial alternating current stimulation entrains alpha oscillations by preferential phase synchronization of fast-spiking cortical neurons to stimulation waveform. Nat Commun, 12(1), 3151.

Hubel, D. H., & Wiesel, T. N. (1968). Receptive fields and functional architecture of monkey striate cortex. J Physiol, 195(1), 215–243.

Jehee, J. F., Ling, S., Swisher, J. D., van Bergen, R. S., & Tong, F. (2012). Perceptual learning selectively refines orientation representations in early visual cortex. J Neurosci, 32(47), 16747–16753a.

Jensen, O., & Mazaheri, A. (2010). Shaping functional architecture by oscillatory alpha activity: gating by inhibition. Front Hum Neurosci, 4(186), 186.

Jeter, P. E., Dosher, B. A., Petrov, A., & Lu, Z. L. (2009). Task precision at transfer determines specificity of perceptual learning. J Vis, 9(3), 1 1–13.

Jia, K., Zamboni, E., Kemper, V., Rua, C., Goncalves, N. R., Ng, A. K. T., … Kourtzi, Z. (2020). Recurrent Processing Drives Perceptual Plasticity. Curr Biol, 4177–4187.

Johnson, L., Alekseichuk, I., Krieg, J., Doyle, A., Yu, Y., Vitek, J., … Opitz, A. (2020). Dose-dependent effects of transcranial alternating current stimulation on spike timing in awake nonhuman primates. Sci Adv, 6(36), eaaz2747.

Kanai, R., Chaieb, L., Antal, A., Walsh, V., & Paulus, W. (2008). Frequency-dependent electrical stimulation of the visual cortex. Curr Biol, 18(23), 1839–1843.

Kanai, R., Paulus, W., & Walsh, V. (2010). Transcranial alternating current stimulation (tACS) modulates cortical excitability as assessed by TMS-induced phosphene thresholds. Clin Neurophysiol, 121(9), 1551–1554.

Kar, K., Duijnhouwer, J., & Krekelberg, B. (2017). Transcranial Alternating Current Stimulation Attenuates Neuronal Adaptation. J Neurosci, 37(9), 2325–2335.

Kar, K., & Krekelberg, B. (2014). Transcranial alternating current stimulation attenuates visual motion adaptation. J Neurosci, 34(21), 7334–7340.

Kasten, F. H., Dowsett, J., & Herrmann, C. S. (2016). Sustained Aftereffect of alpha-tACS Lasts Up to 70 min after Stimulation. Front Hum Neurosci, 10, 245.

Kasten, F. H., Duecker, K., Maack, M. C., Meiser, A., & Herrmann, C. S. (2019). Integrating electric field modeling and neuroimaging to explain inter-individual variability of tACS effects. Nat Commun, 10(1), 5427.

Kasten, F. H., Maess, B., & Herrmann, C. S. (2018). Facilitated Event-Related Power Modulations during Transcranial Alternating Current Stimulation (tACS) Revealed by Concurrent tACS-MEG. eNeuro, 5(3).

Keller, A. J., Roth, M. M., & Scanziani, M. (2020). Feedback generates a second receptive field in neurons of the visual cortex. Nature, 582(7813), 545–549.

Klimesch, W., Sauseng, P., & Hanslmayr, S. (2007). EEG alpha oscillations: the inhibition-timing hypothesis. Brain Res Rev, 53(1), 63–88.

Krause, M. R., Vieira, P. G., Csorba, B. A., Pilly, P. K., & Pack, C. C. (2019). Transcranial alternating current stimulation entrains single-neuron activity in the primate brain. Proc Natl Acad Sci U S A, 116(12), 5747–5755.

La Rocca, D., Ciuciu, P., Engemann, D. A., & van Wassenhove, V. (2020). Emergence of beta and gamma network following multisensory training. Neuroimage, 206, 116313.

Lange, J., Oostenveld, R., & Fries, P. (2013). Reduced occipital alpha power indexes enhanced excitability rather than improved visual perception. J Neurosci, 33(7), 3212–3220.

Law, C.-T., & Gold, J. I. (2008). Neural correlates of perceptual learning in a sensory-motor, but not a sensory, cortical area. Nat Neurosci, 11(4), 505–513.

Levi, D. M., & Li, R. W. (2009). Improving the performance of the amblyopic visual system. Philos Trans R Soc Lond B Biol Sci, 364(1515), 399–407.

Liu, A., Voroslakos, M., Kronberg, G., Henin, S., Krause, M. R., Huang, Y., … Buzsaki, G. (2018). Immediate neurophysiological effects of transcranial electrical stimulation. Nat Commun, 9(1), 5092.

Loewenstein, Y., Raviv, O., & Ahissar, M. (2021). Dissecting the Roles of Supervised and Unsupervised Learning in Perceptual Discrimination Judgments. J Neurosci, 41(4), 757–765.

Lu, J., Luo, L., Wang, Q., Fang, F., & Chen, N. (2021). Cue-triggered activity replay in human early visual cortex. Sci China Life Sci, 64(1), 144–151.

Maniglia, M., & Seitz, A. R. (2018). Towards a whole brain model of Perceptual Learning. Curr Opin Behav Sci, 20, 47–55.

Mathewson, K. E., Beck, D. M., Ro, T., Maclin, E. L., Low, K. A., Fabiani, M., & Gratton, G. (2014). Dynamics of alpha control: preparatory suppression of posterior alpha oscillations by frontal modulators revealed with combined EEG and event-related optical signal. J Cogn Neurosci, 26(10), 2400–2415.

Mayhew, S. D., Li, S., & Kourtzi, Z. (2012). Learning acts on distinct processes for visual form perception in the human brain. J Neurosci, 32(3), 775–786.

Mejias, J. F., Murray, J. D., Kennedy, H., & Wang, X. J. (2016). Feedforward and feedback frequency-dependent interactions in a large-scale laminar network of the primate cortex. Sci Adv, 2(11), e1601335.

Michalareas, G., Vezoli, J., van Pelt, S., Schoffelen, J. M., Kennedy, H., & Fries, P. (2016). Alpha-Beta and Gamma Rhythms Subserve Feedback and Feedforward Influences among Human Visual Cortical Areas. Neuron, 89(2), 384–397.

Moldakarimov, S., Bazhenov, M., & Sejnowski, T. J. (2014). Top-down inputs enhance orientation selectivity in neurons of the primary visual cortex during perceptual learning. PLoS Comput Biol, 10(8), e1003770.

Mukai, I., Kim, D., Fukunaga, M., Japee, S., Marrett, S., & Ungerleider, L. G. (2007). Activations in visual and attention-related areas predict and correlate with the degree of perceptual learning. J Neurosci, 27(42), 11401–11411.

Muller-Gass, A., Duncan, M., & Campbell, K. (2017). Brain states predict individual differences in perceptual learning. Pers Individ Dif, 118, 29–38.

Nakazono, H., Ogata, K., Takeda, A., Yamada, E., Kimura, T., & Tobimatsu, S. (2020). Transcranial alternating current stimulation of α but not β frequency sharpens multiple visual functions. Brain Stimul, 13(2), 343–352.

Neuling, T., Rach, S., & Herrmann, C. S. (2013). Orchestrating neuronal networks: sustained after-effects of transcranial alternating current stimulation depend upon brain states. Front Hum Neurosci, 7, 161.

Nikolaev, A. R., Gepshtein, S., & van Leeuwen, C. (2016). Intermittent regime of brain activity at the early, bias-guided stage of perceptual learning. J Vis, 16(14).

Opitz, A., Falchier, A., Yan, C. G., Yeagle, E. M., Linn, G. S., Megevand, P., … Schroeder, C. E. (2016). Spatiotemporal structure of intracranial electric fields induced by transcranial electric stimulation in humans and nonhuman primates. Sci Rep, 6, 31236.

Pelli, D. G. (1997). The VideoToolbox software for visual psychophysics: Transforming numbers into movies. Spat Vis, 10(4), 437–442.

Pogosyan, A., Gaynor, L. D., Eusebio, A., & Brown, P. (2009). Boosting cortical activity at Beta-band frequencies slows movement in humans. Curr Biol, 19(19), 1637–1641.

R Core Team. (2020). R: A Language and Environment for Statistical Computing (Version 4.0.3). Vienna, Austria: R Foundation for Statistical Computing. Retrieved from http://www.R-project.org/

Reinhart, R. M. G., & Nguyen, J. A. (2019). Working memory revived in older adults by synchronizing rhythmic brain circuits. Nat Neurosci, 22(5), 820–827.

Roberts, D. M., Fedota, J. R., Buzzell, G. A., Parasuraman, R., & McDonald, C. G. (2014). Prestimulus oscillations in the alpha band of the EEG are modulated by the difficulty of feature discrimination and predict activation of a sensory discrimination process. J Cogn Neurosci, 26(8), 1615–1628.

Rodan, A., Candela Marroquin, E., & Jara Garcia, L. C. (2020). An updated review about perceptual learning as a treatment for amblyopia. J Optom.

Romei, V., Brodbeck, V., Michel, C., Amedi, A., Pascual-Leone, A., & Thut, G. (2008). Spontaneous fluctuations in posterior alpha-band EEG activity reflect variability in excitability of human visual areas. Cereb Cortex, 18(9), 2010–2018.

Sanayei, M., Chen, X., Chicharro, D., Distler, C., Panzeri, S., & Thiele, A. (2018). Perceptual learning of fine contrast discrimination changes neuronal tuning and population coding in macaque V4. Nat Commun, 9(1), 4238.

Schafer, R., Vasilaki, E., & Senn, W. (2007). Perceptual learning via modification of cortical top-down signals. PLoS Comput Biol, 3(8), e165.

Schnitzler, A., & Gross, J. (2005). Normal and pathological oscillatory communication in the brain. Nat Rev Neurosci, 6(4), 285–296.

Schoups, A., Vogels, R., Qian, N., & Orban, G. (2001). Practising orientation identification improves orientation coding in V1 neurons. Nature, 412(6846), 549–553.

Shibata, K., Sasaki, Y., Bang, J. W., Walsh, E. G., Machizawa, M. G., Tamaki, M., … Watanabe, T. (2017). Overlearning hyperstabilizes a skill by rapidly making neurochemical processing inhibitory-dominant. Nat Neurosci, 20(3), 470–475.

Sigala, R., Haufe, S., Roy, D., Dinse, H. R., & Ritter, P. (2014). The role of alpha-rhythm states in perceptual learning: insights from experiments and computational models. Front Comput Neurosci, 8, 36.

Struber, D., Rach, S., Neuling, T., & Herrmann, C. S. (2015). On the possible role of stimulation duration for after-effects of transcranial alternating current stimulation. Front Cell Neurosci, 9, 311.

Struber, D., Rach, S., Trautmann-Lengsfeld, S. A., Engel, A. K., & Herrmann, C. S. (2014). Antiphasic 40 Hz oscillatory current stimulation affects bistable motion perception. Brain Topogr, 27(1), 158–171.

Theves, S., Chan, J. S., Naumer, M. J., & Kaiser, J. (2020). Improving audio-visual temporal perception through training enhances beta-band activity. Neuroimage, 206, 116312.

Thut, G., Miniussi, C., & Gross, J. (2012). The functional importance of rhythmic activity in the brain. Curr Biol, 22(16), R658–663.

Toosi, T. E K. T., & Esteky, H. (2017). Learning temporal context shapes prestimulus alpha oscillations and improves visual discrimination performance. J Neurophysiol, 118(2), 771–777.

Tsodyks, M., & Gilbert, C. (2004). Neural networks and perceptual learning. Nature, 431(7010), 775–781.

Turi, Z., Ambrus, G. G., Janacsek, K., Emmert, K., Hahn, L., Paulus, W., & Antal, A. (2013). Both the cutaneous sensation and phosphene perception are modulated in a frequency-specific manner during transcranial alternating current stimulation. Restor Neurol Neurosci, 31(3), 275–285.

van Dijk, H., Schoffelen, J. M., Oostenveld, R., & Jensen, O. (2008). Prestimulus oscillatory activity in the alpha band predicts visual discrimination ability. J Neurosci, 28(8), 1816–1823.

van Kerkoerle, T., Self, M. W., Dagnino, B., Gariel-Mathis, M. A., Poort, J., van der Togt, C., & Roelfsema, P. R. (2014). Alpha and gamma oscillations characterize feedback and feedforward processing in monkey visual cortex. Proc Natl Acad Sci U S A, 111(40), 14332–14341.

Van Meel, C., Daniels, N., de Beeck, H. O., & Baeck, A. (2016). Effect of tDCS on task relevant and irrelevant perceptual learning of complex objects. J Vis, 16(6), 1–15.

VanRullen, R. (2016). Perceptual Cycles. Trends Cogn Sci, 20(10), 723–735.

Vieira, P. G., Krause, M. R., & Pack, C. C. (2020). tACS entrains neural activity while somatosensory input is blocked. PLoS Biol, 18(10), e3000834.

Voroslakos, M., Takeuchi, Y., Brinyiczki, K., Zombori, T., Oliva, A., Fernandez-Ruiz, A., … Berenyi, A. (2018). Direct effects of transcranial electric stimulation on brain circuits in rats and humans. Nat Commun, 9(1), 483.

Vossen, A., Gross, J., & Thut, G. (2015). Alpha Power Increase After Transcranial Alternating Current Stimulation at Alpha Frequency (α-tACS) Reflects Plastic Changes Rather Than Entrainment. Brain Stimul, 8(3), 499–508.

Wach, C., Krause, V., Moliadze, V., Paulus, W., Schnitzler, A., & Pollok, B. (2013). Effects of 10 Hz and 20 Hz transcranial alternating current stimulation (tACS) on motor functions and motor cortical excitability. Behav Brain Res, 241(0), 1–6.

Watanabe, T., & Sasaki, Y. (2015). Perceptual learning: toward a comprehensive theory. Annu Rev Psychol, 66, 197–221.

Watson, A. B., & Pelli, D. G. (1983). QUEST: a Bayesian adaptive psychometric method. Percept Psychophys, 33(2), 113–120.

Weiss, Y., Edelman, S., & Fahle, M. (1993). Models of Perceptual Learning in Vernier Hyperacuity. Neural Computation, 5(5), 695–718.

Yan, Y., Rasch, M. J., Chen, M., Xiang, X., Huang, M., Wu, S., & Li, W. (2014). Perceptual training continuously refines neuronal population codes in primary visual cortex. Nat Neurosci, 17(10), 1380–1387.

Yan, Y., Zhaoping, L., & Li, W. (2018). Bottom-up saliency and top-down learning in the primary visual cortex of monkeys. Proc Natl Acad Sci U S A, 115(41), 10499–10504.

Yang, T., & Maunsell, J. H. (2004). The effect of perceptual learning on neuronal responses in monkey visual area V4. J Neurosci, 24(7), 1617–1626.

Yu, Q., Zhang, P., Qiu, J., & Fang, F. (2016). Perceptual Learning of Contrast Detection in the Human Lateral Geniculate Nucleus. Curr Biol, 26(23), 3176–3182.

Zaehle, T., Rach, S., & Herrmann, C. S. (2010). Transcranial alternating current stimulation enhances individual alpha activity in human EEG. PLoS One, 5(11), e13766.

Zazio, A., Schreiber, M., Miniussi, C., & Bortoletto, M. (2020). Modelling the effects of ongoing alpha activity on visual perception: The oscillation-based probability of response. Neurosci Biobehav Rev, 112, 242–253.

Zhang, P., Hou, F., Yan, F. F., Xi, J., Lin, B. R., Zhao, J., … Huang, C. B. (2018). High reward enhances perceptual learning. J Vis, 18(8), 1–21.

Zhang, Y., Zhang, Y., Cai, P., Luo, H., & Fang, F. (2019). The causal role of alpha-oscillations in feature binding. Proc Natl Acad Sci U S A, 116(34), 17023–17028.

